# Identification of bacterial drug-resistant cells by the convolutional neural network in transmission electron microscope images

**DOI:** 10.1101/2021.10.19.464925

**Authors:** Mitsuko Hayashi-Nishino, Kota Aoki, Akihiro Kishimoto, Yuna Takeuchi, Aiko Fukushima, Kazushi Uchida, Tomio Echigo, Yasushi Yagi, Mika Hirose, Kenji Iwasaki, Eitaro Shin’ya, Takashi Washio, Chikara Furusawa, Kunihiko Nishino

## Abstract

The emergence of bacteria that are resistant to antibiotics is common in areas where antibiotics are used widely. The current standard procedure for detecting bacterial drug resistance is based on bacterial growth under antibiotic treatments. Here we describe the morphological changes in enoxacin-resistant *Escherichia coli* cells and the computational method used to identify these resistant cells in transmission electron microscopy (TEM) images without using antibiotics. Our approach was to create patches from TEM images of enoxacin-sensitive and enoxacin-resistant *E. coli* strains, use a convolutional neural network for patch classification, and identify the strains on the basis of the classification results. The proposed method was highly accurate in classifying cells, achieving an accuracy rate of 0.94. Using a gradient-weighted class activation mapping to visualize the region of interest, enoxacin-resistant and enoxacin-sensitive cells were characterized by comparing differences in the envelope. Moreover, Pearson’s correlation coefficients suggested that four genes, including *lpp*, the gene encoding the major outer membrane lipoprotein, were strongly associated with the image features of enoxacin-resistant cells.

## Introduction

The development of antibiotics has progressed dramatically since the middle of the 20^th^ century, but drug-resistant bacteria emerged shortly after antibacterial drugs were introduced to treat bacterial infections, and antibiotic-resistant strains have increased rapidly in number with the long-term overuse of antibacterial drugs. Consequently, in recent years, the emergence of multidrug-resistant bacteria with resistance to multiple types of antibiotics has become a global problem. Various types of resistant bacteria have appeared in clinic, and effective countermeasures to combat multidrug-resistant bacteria are required.

Laboratory-based bacterial evolution is a powerful tool for investigating the dynamics of acquired drug resistance.^1, 2, 3^ In these experiments, bacterial cells are exposed to fixed concentrations of drugs, around which cell growth is partially or completely inhibited such that a selective advantage for resistant strains is maintained. Recently, Suzuki *et al*. performed laboratory evolution of *Escherichia coli* under various drug treatment conditions to obtain resistant strains, including those resistant to quinolones such as enoxacin (ENX).^1^ For each drug-resistant strain obtained, transcriptome and genome sequencing analyses were performed to identify fixed mutations and gene expression changes. By integrating these data and using a simple mathematical model, they succeeded in quantitatively predicting resistance to various drugs on the basis of the gene expression levels.^1^ Because many gene expression changes were observed in the drug-resistant strains, we queried whether these changes may affect the bacterial morphology. There are several reports on the effect of antibiotics on bacterial morphology, including morphogenesis and fatal variations of staphylococci in the presence of penicillin^4^ and the response of *E. coli* to quinolones;^5^ however, there is little known about the morphology of drug-resistant bacteria compared with that of the drug-sensitive parent strain, especially in the absence of drugs.

In this study, we performed a morphological analysis of ENX-resistant cells that were obtained through the laboratory evolution of *E. coli*. Microscope images of both drug-resistant and drug-sensitive cells were obtained and their morphological characteristics were described. Electron microscopy, including transmission electron microscopy (TEM), is a powerful means of observing the ultrastructures of various biological samples, and these tools have been used often to study bacterial cell morphology. Recently, it was reported that deep learning approaches have been applied to electron microscopy images of biological specimens, including isolated macromolecules and brain tissues.^6, 7^ However, to date, few computational methods have been developed to identify drug-resistant bacteria in TEM images. Considering this background, clarifying the morphological changes that occur in bacterial drug resistance is important, and the ability to estimate drug resistance from the morphological features of bacterial cells is also key to basic and applied microbiological studies. We therefore attempted to distinguish drug-resistant cells according to their morphological features visible on TEM images using deep learning and identify the discriminatory features. Moreover, we sought to identify the genes related to the morphological characteristics of ENX-resistant cells.

This study provides a novel method for discriminating drug-resistant and drug-sensitive bacterial cells from their TEM images using deep learning. The main contributions of this paper include the 1) clarification of morphological changes in drug-resistant bacteria from observations using optical and electron microscopy, 2) application of deep learning to discriminate TEM images of drug-resistant bacterial cells with a high level of accuracy, 3) visualization of morphological features peculiar to the drug-resistant bacteria using classifier models, and 4) identification of genes that may be associated with the morphological changes by correlating the image features extracted by the classifier models with gene expression levels.

## Results

### Observation of drug-resistant and drug-sensitive bacterial strains using optical microscopy

The morphologies of both drug-resistant *E. coli* cells and sensitive parental cells were observed first under a light microscope. The parental *E*.*coli* strain and drug-resistant laboratory-evolved strains obtained from a previous study^1^ were grown in the absence of antibiotics. Of 10 different drug-resistant strains, four types of strain with high resistance to drugs with different mechanisms of action [ENX, amikacin (AMK), cefixime (CFIX), and chloramphenicol (CP)] were selected for observation. Although other drug-resistant strains exhibited rod-shaped cell morphology, the ENX-resistant strains were more spherical in cell shape (Fig. 1A). Further morphometric analysis of the ENX-resistant strains demonstrated that the distribution of major and minor axis lengths appeared to have shifted from those of the parental strain (Fig. 1B). Although the cells of the ENX-resistant strains tended to be shorter, they were wider. Consequently, the aspect ratio was smaller than that of the parental strain (Fig. 1C). Furthermore, the aspect ratio of the ENX-resistant strains was significantly lower than that for the other drug-resistant strains, strongly suggesting that the cell shape of the ENX strains had changed from that of the parental strain (Fig. 1D). Conversely, the aspect ratio of the AMK-resistant strains was larger than that of the parental strain.

**Fig. 1.**
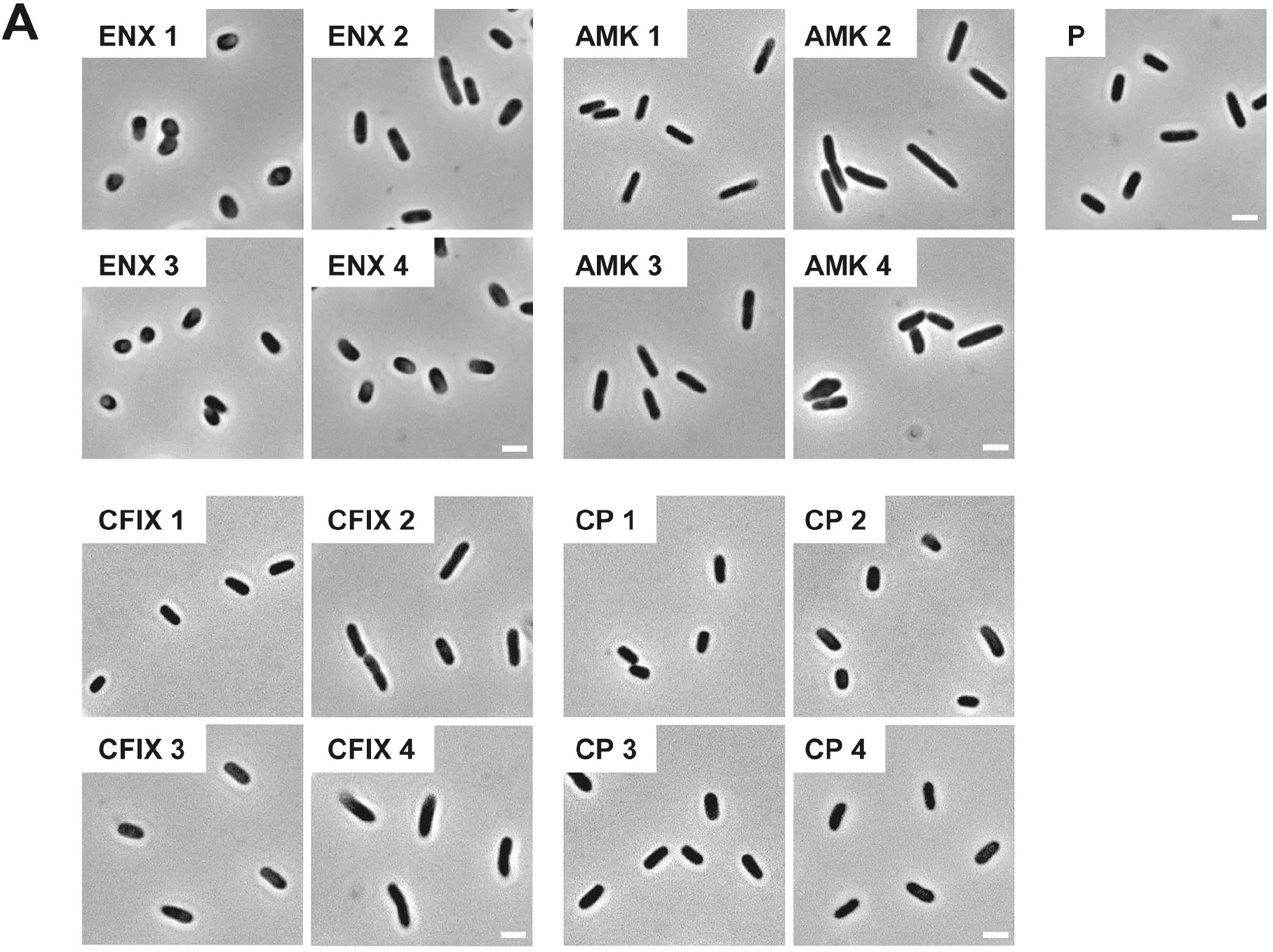
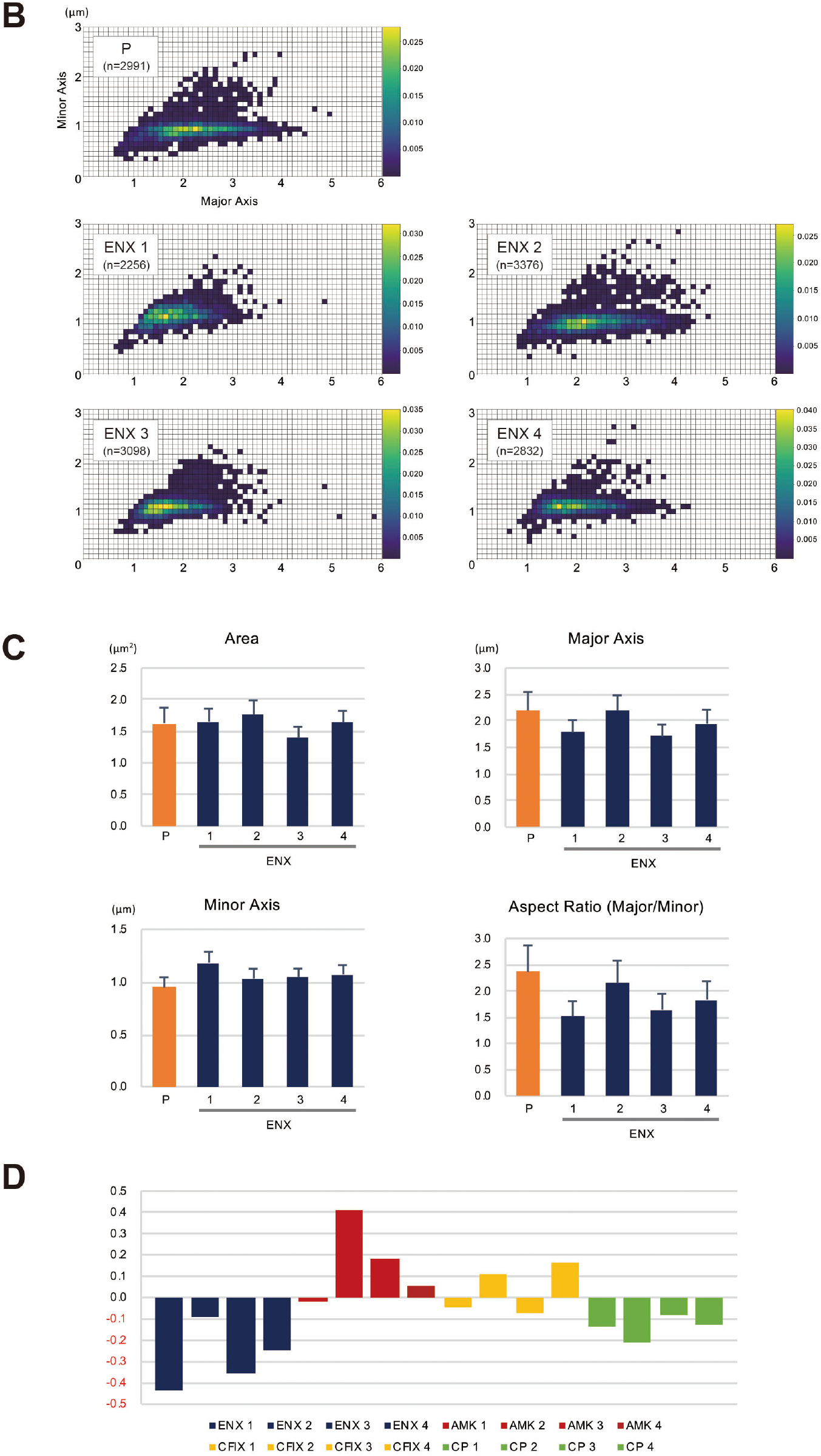
Light microscopy of drug-resistant strains. (A) Representative images of four different drug-resistant strains, enoxacin (ENX), amikacin (AMK), cefixime (CFIX), chloramphenicol (CP), and the parental strain (P) are shown. Scale bars, 2 μm. (B–D) Results of the morphometric analysis presented as (B) scatter plots of major vs. minor axis lengths of the parental cells and ENX-resistant cells. The color scales indicate the ratio of cell densities. (C) Bar graphs show averages of bacterial cell sizes, major and minor axis lengths, and the aspect ratio. All indicated data were significantly different by *P* < 0.01, Student’s *t* test. (D) Logarithmic changes in the average aspect ratio of the indicated drug-resistant strains against the parental strain are shown.

### Observation of ENX-resistant strains using TEM

The results of the light microscopy study suggested that the ENX-resistant strains exhibited the largest difference in cell shape of all the drug-resistant strains investigated. We therefore speculated that the ultrastructures of these strains might also be different from those of the parental strain. Hence, we used TEM to further study the detailed morphologies of the ENX-resistant strains. A high-pressure freezing procedure was employed to fix the samples and preserve the ultrastructures of the bacterial cells better than when classical chemical fixation is used. ^8, 9^ Both the ENX-resistant strains and the parental strain cultured in modified M9 medium were cryofixed and freeze-substituted. The bacterial specimens were further processed with thin-sectioning and staining as illustrated in Figure S1. Finally, the specimens were observed using TEM and their representative images are depicted in Figure 2. Cross-sections of bacterial cells were included in each image, and the envelope including the outer membrane and periplasm, granular structures, and cytoplasm were visible. Occasionally, the periplasmic space appeared to be enlarged around the cell poles in both the parental and ENX-resistant cells, except ENX 4 cells (Fig. 2, lower panels). This enlargement of the periplasm was considered partially to be the result of the cryoprocedures as reported previously,^10^ although the degree of enlargement appeared to be different among the strains. In the parental cells, dense granules appeared to have accumulated at the cell poles, whereas they seemed less electron dense and relatively less abundant in the ENX-resistant cells. When the morphologies of four ENX-resistant strains were observed, the appearance of the cell shape, the outer membrane, periplasmic space, and granules varied (Fig. 2, upper panels). Although a rod-like cell shape was seen in the parental strain, the cell shape appeared to be irregular and relatively round in the ENX 1, ENX 3, and ENX 4 strains; this difference was reflected in the results obtained using light microscopy. Additionally, the outer membranes of the ENX-resistant cells appeared to be slightly wavy, and bleb-like structures were occasionally visible, especially in the ENX 3 and ENX 4 cells (Fig. 2, lower panels). Taken together, these results suggested that the ENX-resistant strains harbored different morphologies from the parental strain at the ultrastructural level.

**Fig. 2.**
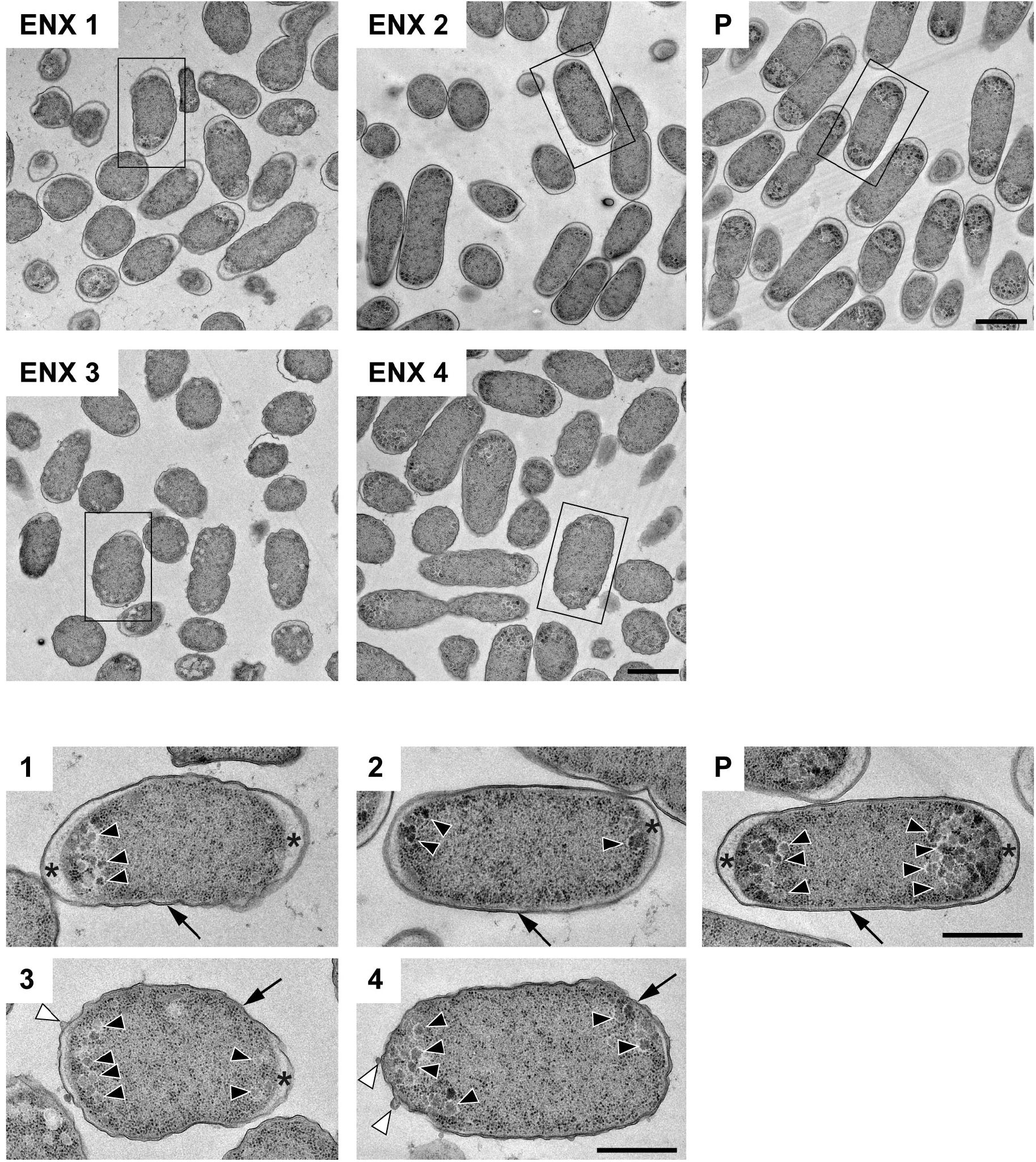
Ultrastructure of the ENX-resistant cells. Upper panels show representative transmission electron micrographs of the ENX-resistant cells and the parental (P) cells. Lower panels show magnification of the boxed bacterial cells in upper panels, depicting the detailed morphology of the cells. The number denotes the indicated ENX-resistant strains. The outer membrane, granules, bleb-like structures, and periplasmic space are indicated by an arrow, arrowheads, white arrowheads, and asterisks, respectively. Scale bars, 1 μm in upper panels; 500 nm in lower panels.

We next examined whether the ENX-resistant strains could be discriminated from the parental strain by machine learning.

### Classification of ENX-resistant strains using convolutional neural networks

The deep learning approach^11^ was reported to achieve much higher levels of performance than conventional, handcrafted feature-based techniques in an image recognition competition (the ImageNet Large Scale Visual Recognition Challenge^12^) held in 2012. Since then, deep learning approaches have been applied to EM images. The DeepEM algorithm^6^ was proposed to determine whether a single particle is present in a boxed area cropped from cryo-EM images for the 3D reconstruction of the structure. The DeepEM3D algorithm^7^ and other methods^13,14^ were proposed for neuronal membrane segmentation in 3D EM image stacks. Furthermore, a cloud-based solution to the segmentation tasks, named CDeep3M,^15^ was developed as an applicable tool for the biomedical community. Some state-of-the-art deep learning models were used to classify scanning electron microscopy images in 10 categories^16^ and to discriminate whether an image was obtained by TEM or scanning TEM.^17^

In the present study, we aimed to discriminate between the ENX-resistant strains and the parental strain, rather than to distinguish an individual strain, and find morphological features shared by the ENX-resistant strains. Our experiments in image classification using convolutional neural network (CNN) models with TEM data sets were conducted as illustrated in Figure 3 to demonstrate differences in the appearance and shape of ENX-resistant strains compared with the parental strain. Because 2048 × 2048 pixel TEM images were too large to be fed into the CNN models, and there were not enough images to train the CNN models effectively, patches that contained portions of bacterial cells were cropped from each image. To validate the consistency and robustness of the CNN models against the process of acquiring the TEM data sets, three blocks for each ENX-resistant strain and six blocks for the parental strain were prepared, respectively, and split into three individual sets for three-fold cross-validation. The TEM images for all of the bacterial specimen blocks used in the experiments showed reproducibility, but there were variations in image quality for each cryofixation block among the strains (Fig. S2A). Although the overall image features appeared to be similar for each strain in different cryofixation blocks, when observed in detail, the intensity of the staining, the appearance of the outer membrane and periplasmic space, and the electron densities of granules varied slightly for each block. Additionally, variations in image quality and cell density were observed among grids, as well as in different regions of the grid sections. Moreover, some pictures contained noncellular structures such as knife marks, a wrinkle in a section, or a small contaminant, which had to be eliminated as much as possible to avoid any influence on the classification result (Fig. S2B). As the TEM images contained cross-sections of cells at different angles, large variations in the appearance of cell shape were observed (Fig. S2C). Therefore, because of these issues, we considered it important to evaluate the robustness of the image data by preparing a multitude of samples for each strain and using image data sets obtained from a multitude of grids.

**Fig. 3.**
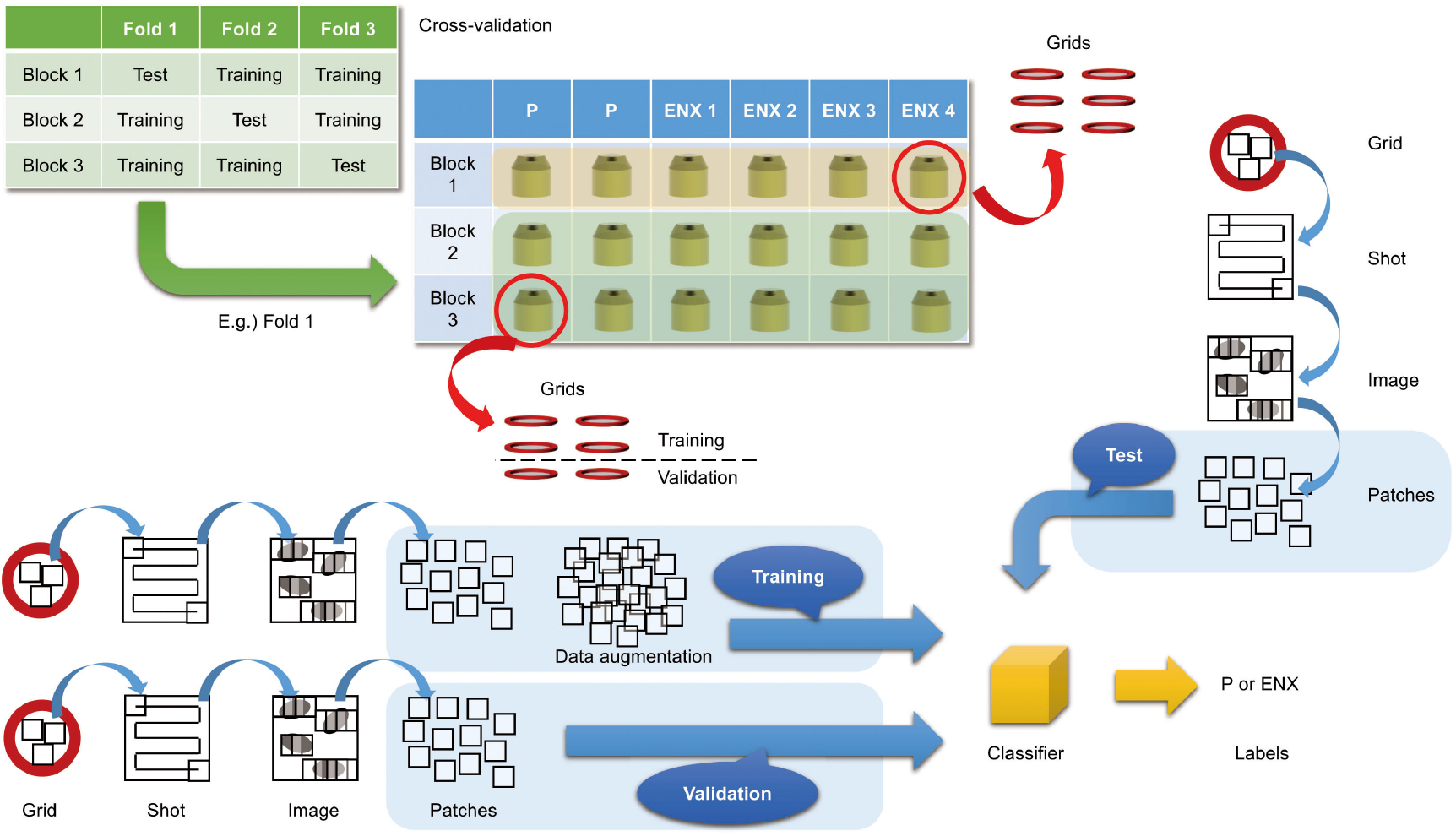
Overview of three-fold cross-validation for prediction of ENX-resistant cells. The cartoon shows that TEM images were taken (denoted as “Shot”) from the 80-nm-thick sections on multiple grids, which were prepared from three blocks for each ENX-resistant strain and six blocks for the parental strain, and were split into three individual sets for three-fold cross-validation. Numerous patches containing portions of cross-sections of bacterial cells were extracted from TEM images and used for training, validation, and tests for classifier models (see also Methods).

The CNN architecture and hyperparameters used in this study are illustrated in Figure 4. The AlexNet model^11^ was adopted because it was found to be efficient and effective at demonstrating the feasibility of TEM image classification. A batch normalized version^18^ of AlexNet was used for patch classifications. After the TEM images were preprocessed and patch data sets constructed (Fig. S3 and see also Methods), the model was trained using the ImageNet dataset^12^ during a pretraining phase and then retrained or fine-tuned to classify the TEM images of the ENX-resistant strains and parental strain. The classifier models were successfully trained, but not overfitted, because the validation losses converged to values that resembled the training losses during every fold of the cross-validation (Fig. 5). Consequently, similar performances were achieved for sensitivity, specificity, and accuracy in the validation sets during all folds.

**Fig. 4.**
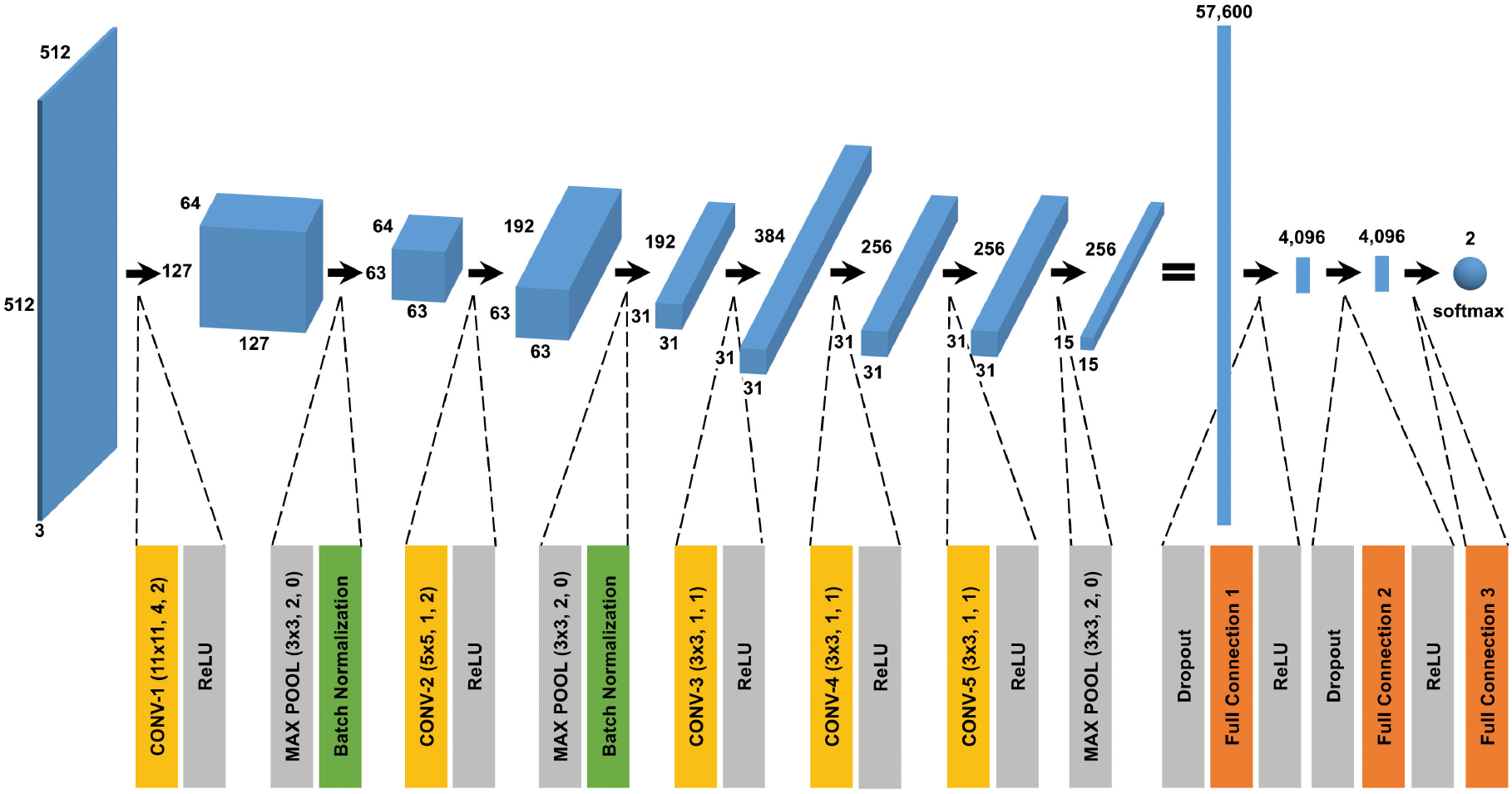
CNN architecture used in this study. A batch normalized version of AlexNet was used for patch classification. The network architecture and some hyperparameters are illustrated in the figure. The softmax function was used for the classification output.

**Fig. 5.**
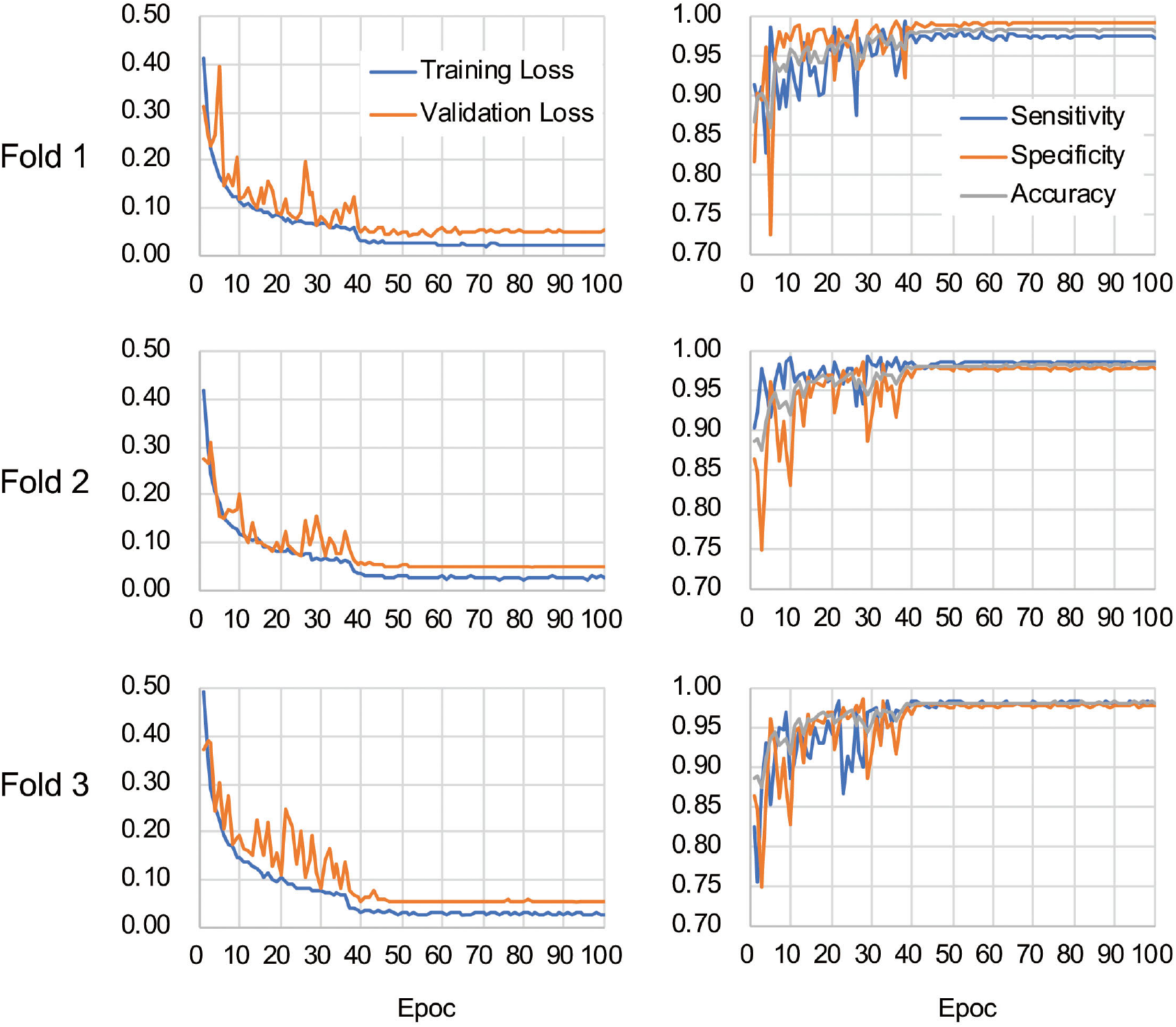
Validation loss and accuracy curves. The history of losses on the training and validation sets during each training phase (left panels) and the history of the sensitivity, specificity, and accuracy scores (right panels) are shown for each fold.

Table 1 shows the performance of the classifier models evaluated in the test sets during the cross-validation scheme. All of the sensitivity and specificity, defined as Eq. (1) and (2), respectively (see Methods), exceeded ∼0.9. The classifier models achieved high levels of performance for TEM patch classifications in the test sets, whereas the sensitivity and specificity scores varied among the folds of the cross-validation. One of the reasons is that the number of patches extracted from the TEM images differed considerably among the bacterial strains (Table S1) and this imbalance in the data may have influenced the training of the classifier models and performance evaluation. Unbalanced data sets were acquired because of the elimination of patches containing noncellular structures and variations in the cell densities among the acquired images, as described above. Moreover, the difference in classification performance may have been caused by variations in the appearance and shape of cells (Fig. S2B and C) that could not be compensated for by increasing the number of training samples/patches. However, overfitting did not seem to occur. That is, the classifier models were successfully trained because there was only a small gap between the losses in the training and validation sets.

**Table 1.** Classification results^a^.

| Fold | True<br>Positive | False<br>Positive | True<br>Negative | False<br>Negative | Sensitivity | Specificity | Accuracy |
| --- | --- | --- | --- | --- | --- | --- | --- |
| 1 | 228,374 | 12,850 | 115,315 | 8,100 | 0.966 | 0.900 | 0.943 |
| 2 | 168,712 | 2,555 | 154,633 | 13,229 | 0.927 | 0.984 | 0.953 |
| 3 | 174,530 | 14,106 | 129,182 | 2,859 | 0.984 | 0.902 | 0.947 |

<sup>a</sup> The table presents both the number of classified patches (left four columns) and classification accuracy rates (right three columns). “True Positive,” False Positive,” “True Negative,” and “False Negative” denote correctly classified ENX-resistant cell patches, parental cell patches classified into the ENX-resistant cell patches, correctly classified parental cell patches, and ENX-resistant cell patches classified into the parental cell patches, respectively. “Sensitivity” and “Specificity” denote the accuracy rate for classifying ENX-resistant cells and accuracy rate for parental cells, respectively.

### Visualization of the discriminative features of ENX-resistant cells

The next stage of the experiment was to clarify the morphological features of ENX-resistant cells using gradient-weighted class activation mapping (Grad-CAM) to visualize important regions in the images, which a CNN model relies on to predict class labels.^19^ We hypothesized that the activated regions might contribute to the ability to discriminate between the bacterial strains and therefore could indicate the characteristic features of bacterial cells.

Figure 6 shows the representative results acquired using Grad-CAM, which were obtained from one of three folds, by accumulating patchwise results, in which the classification confidence was equal to 1 on the corresponding images. All folds demonstrated similar results (Fig. S4). The classifier models were revealed to be activated predominantly at the envelope for all ENX-resistant strains. Some bleb-like structures on the outer membrane were also activated in both ENX 3 and ENX 4 strains. Thus, these results supported our findings from the initial TEM observation. Besides the envelope, the granules and a portion of cytoplasm in some of the ENX-resistant cells (Fig. 6 and Fig. S4, insets) were occasionally activated. However, the classifier models focused specifically on the dense granules at the cell poles in the parent strain.

**Fig. 6.**
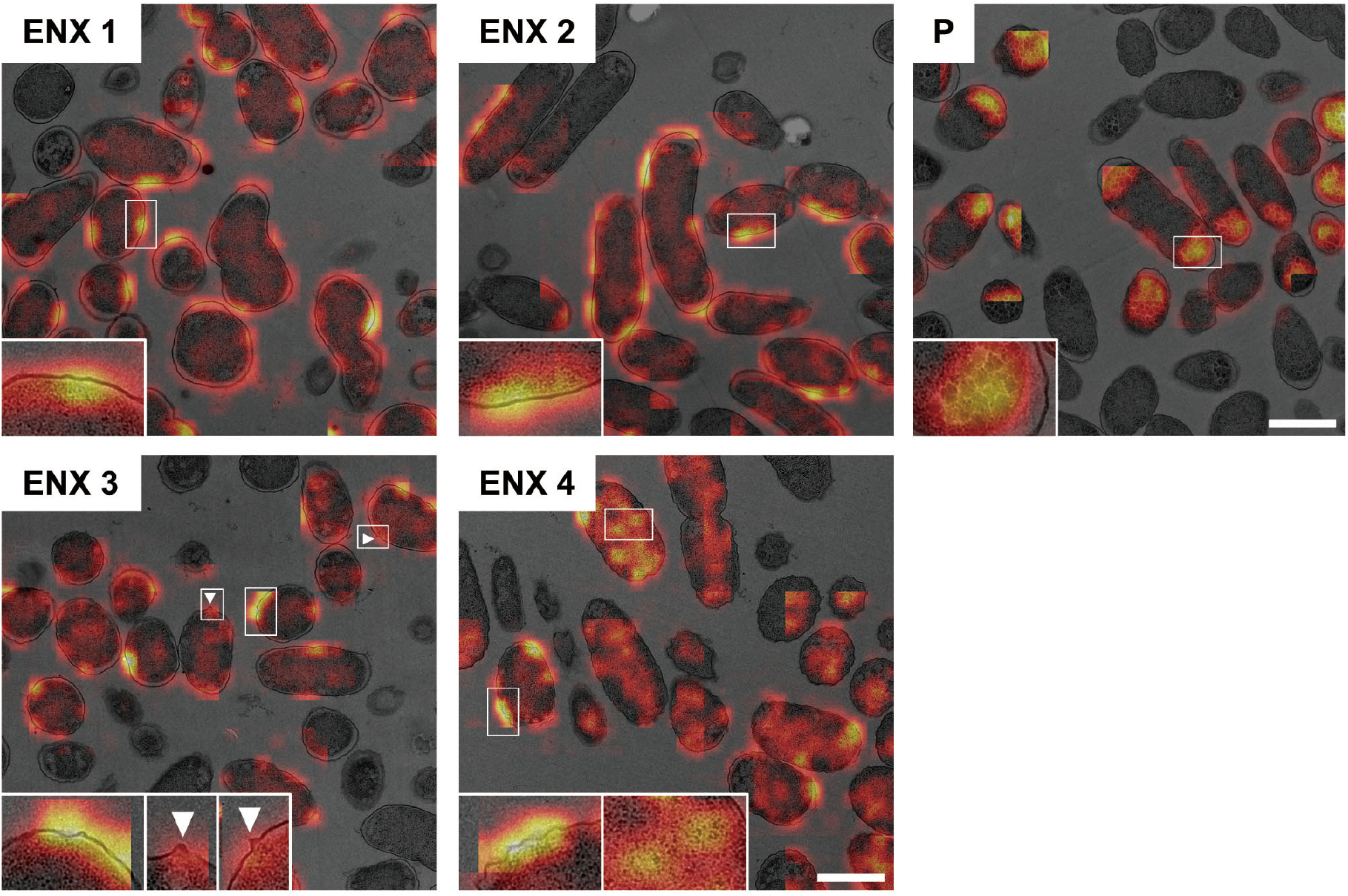
Visualization of the discriminative parts of bacterial cells using Grad-CAM. The representative Grad-CAM images generated from fold three by the accumulation of patchwise results on their corresponding images are shown. Insets depict the magnification of the boxed regions in each image. White arrowheads indicate bleb-like structures. Scale bars, 1 μm.

### Correlation of image features with gene expression

The results obtained using Grad-CAM suggested that the morphological features of the ENX-resistant strains could be found mainly at the envelope, and this led us to consider which genes might be associated with the morphological characteristics of these strains. Thus, we calculated Pearson’s correlation coefficients for gene expression^1^ and the image features extracted from the CNN model [Eq. (3) in Methods]. We retained pairs of genes and image features meeting the condition that the absolute correlation coefficient between them was equal to or greater than 0.999. Consequently, multiple genes were identified for each fold (Table S2) and four appeared in all the folds (Table 2). Surprisingly, all of these four genes were associated with envelope, being either envelope components or involved in the regulation of membrane lipid A modification.^20,21,22,23^ Particularly, *lpp*, a gene that encodes the major outer membrane lipoprotein,^24^ appeared most frequently in all the folds. Lpp is important to the maintenance of the outer membrane structure and its mutant causes morphological alterations in the membrane, resulting in blebs and wavy appearance.^25,26^ Of note, similar morphological features were observed at the outer membrane in both ENX 3-and ENX 4-resistant strains (Fig. 2). In fact, *lpp* in these strains caused a dysfunctional mutation and a significant decrease in gene expression.^1^ Some other genes highly associated with certain image features were identified statistically, but it was difficult to ascertain which gene was associated with each image feature, so these questions remain unanswered.

**Table 2.**
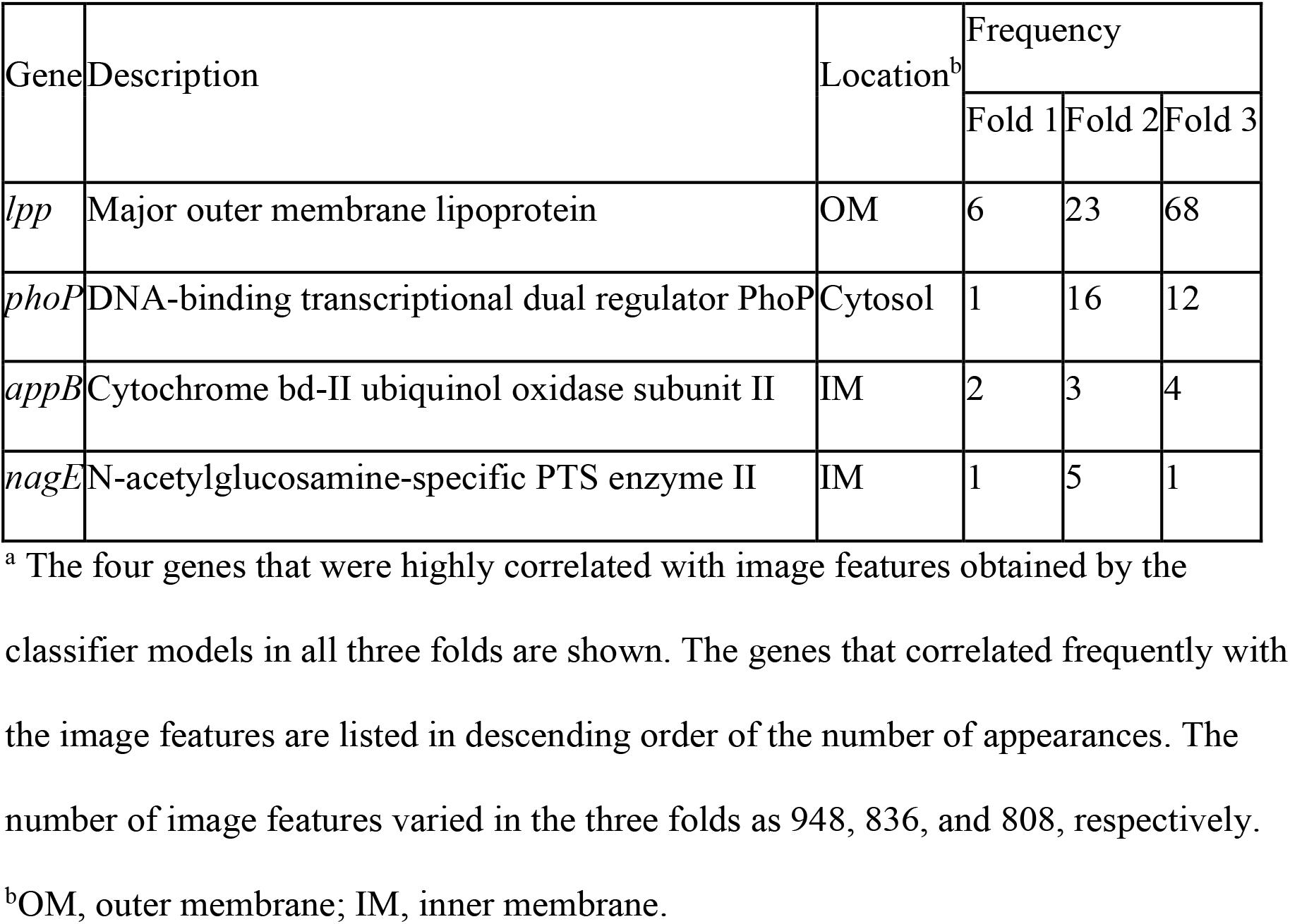
Genes exhibiting a high correlation with image features^a^.

| Gene | Description | Location<sup>b</sup> | Frequency |  |  |
| --- | --- | --- | --- | --- | --- |
|  |  |  | Fold 1 | Fold 2 | Fold 3 |
| *lpp* | Major outer membrane lipoprotein | OM | 6 | 23 | 68 |
| *phoP* | DNA-binding transcriptional dual regulator PhoP | Cytosol | 1 | 16 | 12 |
| *appB* | Cytochrome bd-II ubiquinol oxidase subunit II | IM | 2 | 3 | 4 |
| *nagE* | N-acetylglucosamine-specific PTS enzyme II | IM | 1 | 5 | 1 |

<sup>a</sup> The four genes that were highly correlated with image features obtained by the classifier models in all three folds are shown. The genes that correlated frequently with the image features are listed in descending order of the number of appearances. The number of image features varied in the three folds as 948, 836, and 808, respectively.
<sup>b</sup>OM, outer membrane; IM, inner membrane.

## Discussion

Previous studies on bacterial drug resistance have generally provided an explanation of the genetic mutations that have occurred in drug-resistant cells. In recent years, the analysis of transcriptional data has revealed that the expression patterns of many genes are altered in drug-resistant cells, and it is becoming possible to predict drug-resistant bacteria from gene expression profiles^1^. Similarly, morphological changes associated 1:1 with a particular gene have been described.^27,28^ However, as mentioned above, the expression patterns of multiple genes are altered in drug-resistant strains, and these changes may have a complex effect on the morphology of the bacterial cells.

In this study, we revealed that drug-resistant strains that evolved in the laboratory maintained their morphological changes even in the absence of drugs. This finding suggested that the genetic changes that occur during the acquisition of drug resistance may induce morphological changes. Using deep learning techniques, which have progressed markedly in recent years, we succeeded in accurately identifying TEM images of strains resistant to ENX, a quinolone antibacterial agent, showing the morphological characteristics of ENX-resistant strains. In the future, it will be possible to explain the mechanisms of multidrug resistance in bacteria in a new way by clarifying the morphological characteristics of various types of drug-resistant strains. Making image discrimination possible for clinical isolates will be a significant advance as well. In this study, patches containing a portion of cells were used to discriminate the resistant strains, and local differences in the internal structure were used for identification purposes. However, both the internal structure and cell shape of drug-resistant bacteria change, and therefore, whether single-cell images can be used for highly accurate identification remains to be investigated.

We further explored the genes associated with the image features of the ENX-resistant strains using the image features extracted from the CNN model. Consequently, genes related to the composition of the envelope, including *lpp*, were found to be highly correlated. These genes coincided with the regions of interest in the images visualized by Grad-CAM, suggesting that genes involved in envelope formation are linked to the characteristics seen in ENX-resistant strains. Interestingly, *lpp, phoP*, and *lolE* are regulated by σ^E^, one of the sigma factors that respond to envelope stress,^29^ and *appB* is regulated by σ^S^, the starvation/stationary phase sigma factor that is also induced by antibiotics.^30,31^ Many of the other genes extracted in this study were found also to be involved in sigma factors and stress responses.^32,33^ Presumably, the stress of continuous exposure to the antibacterial drugs in the evolution experiments may have altered the expression of these genes and consequently affected the morphological changes in the ENX-resistant strains. The genes extracted in this study have not previously been identified in reports on evolved drug-resistant bacterial strains,^1,34^ their potential involvement in the morphological changes of the ENX-resistant strains is of great interest. However, it is not clear at present whether these genes correlate highly with the image features of ENX-resistant strains specifically, and further investigations using other drug-resistant strains are required to clarify this point. Furthermore, the morphology of clinical isolates should be compared with that of experimentally evolved strains using image discrimination and genetic analysis to determine whether the morphology of clinical isolates is similarly altered.

The mechanism underlying bacterial resistance to quinolones is assumed to be associated with mutations in the target factors DNA gyrase and topoisomerase IV, decreased expression of porins, and increased activity of efflux pumps.^35^ However, the genes found to be highly correlated with the image features of the ENX-resistant strains were different from these factors, although genetic mutations and expression changes of these factors occurred in these strains.^1^ Thus, the relationship between the factors involved in drug resistance and the morphological changes that occur during the acquisition of drug resistance should be elucidated in future work.

## Methods

### Cell culture conditions

Laboratory-evolved ENX, AMK, CFIX, and CP-resistant strains and the parental MDS42 strain^1^ were used for the experiments. First, a single colony of these resistant strains, harboring the equivalent MIC value of the original stocks, was obtained and stored in an M9 medium containing 15% glycerol at −80°C. Bacterial cells were precultured in a 200 μL modified M9 medium^36^ in Nunc 96-well microplates (Thermo Fisher Scientific Inc.), shaken at 432 rpm on a multimode microplate reader (Infinite M200 PRO, TECAN Ltd.) at 34°C for 23 h, and used for experiments.

### Light microscopy

For light microscopy-based observations, precultures of ENX-, AMK-, CFIX-, and CP-resistant strains and the parental strain were diluted to an OD_600 nm_ of 1 × 10^−4^ to 1 × 10^−8^ in 5 mL of modified M9 medium in glass test tubes and grown in a water bath at 34°C that was shaking at 150 rpm to reach an OD_600 nm_ in the range 0.07–0.13. Bacterial cultures were then transferred to 15 mL centrifuge tubes, and cells were collected by centrifugation at 3000 × g for 10 min at 4°C. The bacterial cell pellets were resuspended with phosphate-buffered saline (PBS, Sigma-Aldrich) and washed twice by centrifugation. Finally, the cell pellets were resuspended with 200 μL of PBS and 2 μL of the cell suspension were mounted on a glass slide and covered with a 22 × 22 mm glass cover slip (Matsunami, Japan). Specimens were then observed, and images were taken under phase-contrast microscopy with an objective lens of ×100 (Leica Microsystems). Individual bacterial cells were detected using a watershed algorithm, and quantitative analysis was performed using Fiji-ImageJ software. The interquartile ranges of the cell size distributions were used for further statistical analysis.

### Fixation and embedding for TEM

Precultures of both ENX-resistant strains and the parental strain were diluted to an OD_600 nm_ of 1 × 10^−4^ to 1 × 10^−8^ in 5 mL of M9 medium in glass test tubes and grown in a water bath at 34°C that was shaking at 150 rpm to reach an OD_600 nm_ in the range 0.07–0.13. Bacterial cultures were then transferred to 15 mL centrifuge tubes and the cells were collected by centrifugation at 3000 × g for 10 min at 4°C. The bacterial pellets (∼8 μL) were resuspended with 3 μL of M9 medium and transferred to Eppendorf tubes and used for cryofixation. Cell suspensions (1.5 μL) were loaded onto a flat specimen carrier (200 μm deep) and frozen multiple times in a Leica EM PACT2 high-pressure freezer (Leica Microsystems). Frozen specimens were transferred into glass bottles with screw caps containing 2% (wt/vol) osmium tetroxide (OsO_4_) in anhydrous acetone as a substitution medium and were freeze-substituted in an automatic freeze substitution unit (EM AFS2; Leica Microsystems). During these periods, the specimens were kept at −80°C for 72 h, and the temperature was raised at a rate of 5°C/h to −20°C and 4°C and kept at each temperature level for 2 h in the substitution medium. Once the samples reached room temperature, the substitution medium was removed and the samples were washed carefully in anhydrous acetone for 5 min, three times. Samples were then incubated in a 1:1 mixture of acetone and Epon 812 resin mixture (TAAB Laboratories) overnight at room temperature, and then in pure Epon mixture overnight. Samples were placed with the cells face up in flat bottomed capsules (TAAB Laboratories), filled with pure Epon mixture, and cured in an oven for polymerization at temperatures of 35°C for 24 h, 42°C for 24 h, and 60°C for 2 days.

### Sectioning and image acquisition

Specimen carriers in Epon blocks were exposed by removing excess resin using a specimen trimming device (EM TRIM2; Leica Microsystems), followed by hand trimming with a razor blade to allow complete exposure. The specimen carrier was detached using a detaching tool according to the manufacturer’s instructions (Leica Microsystems). Ultrathin (80 nm) sections of the cell specimens were cut using an ultramicrotome (EM UC7; Leica Microsystems) equipped with a diamond knife and were collected on formvar/carbon-coated Maxtaform finder grids (Electron Microscopy Sciences). The grids were stained in 2% (wt/vol) aqueous uranyl acetate for 25 min and in lead citrate for 3 min. Ultrathin sections were observed using JEOL TEM (JEM-2100 HC) at an accelerating voltage of 80 kV, and images were taken with a 2k × 2k CCD camera (UltraScan 1000; Gatan Inc.). For machine learning, raw TEM images were taken using automatic image acquisition software (Shot Meister; System in Frontier Inc) at an xy pixel size of 3.4 nm × 3.4 nm, and a single image size of 6.926 μm × 6.926 μm. A panoramic image (denoted as a “Shot” in Fig. 3) comprising 36 images (41.554 μm × 41.554 μm in size) was taken without any margin between neighboring images. Each image was taken with an exposure time of 1 s, giving a 3 s wait time to minimize the image drift. At least five different shots (a total of 180 images) were taken from different arbitrary regions of the section on an individual grid. Hence, at least 900 images were obtained from five grids prepared from each specimen block, and these were used for the analysis.

### Preparation of data sets for machine learning

To create a dataset that was robust against image data variability, a total of three data sets were created, with one set contained each block for ENX-resistant strains and two blocks for a parental strain, giving a total of six blocks. Since there were four lines of ENX-resistant strains and only one line of the parental strain, we used two blocks of the parental strain for each data set and six blocks in total, which was the maximum number to be prepared from one culture, to eliminate the data imbalance as much as possible. Moreover, to cover the variation in image data caused by thin-sectioning and staining, images were taken from five independent grids for each block, as described above. Using these data sets, we replaced each set with training and testing and performed a three-fold cross-validation to evaluate robustness against variations in specimen preparation (see Fig. 3).

### Image preprocessing and construction of a patch data set

Preprocessing was needed for raw TEM images so that the classifier models could handle them effectively. There were some TEM images whose intensity distribution was observed in a dynamic range different from others, and unusual or defective pixels whose intensity value was extremely small or large. Such inconsistency in the intensity values may prevent a classifier model from learning the essential differences in TEM images between drug-resistant strains and the parental strain and thus reduce the classification performance.

A TEM image obtained originally as 16 bit was standardized and converted to 8 bit as follows. Both a mean and a standard deviation of intensity values were first calculated from all pixels of an image, which formed upper and lower bounds on intensity values in accordance with the so-called three-sigma rule. The image was processed so that the intensity values were within the range and then was standardized as if the intensity values followed a standard normal distribution (with zero mean and unit variance). Finally, the image was converted into 8 bit by scaling the intensity values to a range of 0–255. Defective pixels with extremely small or large intensity values could be corrected after completing these steps. A raw TEM image was also subject to uneven intensity levels. For example, intensity levels slightly declined (and thus pixels looked darker) from left to right or top to bottom in an image, and it was considered that this unevenness in intensity was caused by the uneven irradiation of the electron beam on the fluorescent screen. Thus, contrast limited adaptive histogram equalization (CLAHE) was used to enhance the local contrast of TEM images and alleviate uneven intensity levels (Fig. S3A, B).

The original TEM images were 2,048 pixels both in width and height, which did not fit into the pretrained classifier model and were difficult to transfer to the GPU memory. Furthermore, compared with data sets used in typical image recognition tasks, a relatively smaller number of TEM images could be obtained, because it took time and effort as described above. Therefore, subimages, or 512 × 512 pixel patches showing parts of cells, were extracted from the TEM images. This produced a large data set and allowed us to train the classifier models effectively. All TEM images were processed as follows after CLAHE:

1. The noise level in the TEM images was so high as to affect the outcome of the following processes. A Gaussian smoothing filter was typically applied to each image to reduce noise.

2. The Otsu method is one of the most commonly used image thresholding techniques and was adopted to separate cell regions from the background in this study (Fig. S3C). This traditional method worked sufficiently well for images obtained after preprocessing because the intensity histograms tended to become bimodal.

3. It was often the case that the enlarged periplasmic space appeared as holes, which were sometimes large in size, in cell regions obtained using the Otsu method. To fill the holes as much as possible, most external contours that ideally corresponded to the cell outer membrane were detected, and then, areas bounded by the contours were filled (Fig. S3D, E). At the same time, small regions below the given threshold were rejected as false detection. Mask images refined through these procedures were generated and used for further processing.

4. Patches containing parts of cells were selected if the number of pixels assigned to cell regions within the patch exceeded a predefined threshold (Fig. S3F). A sliding window with a stride of 128 pixels was applied to check whether a patch satisfied the above condition. Consequently, a million patches were obtained from each data set of cross-validation folds.

### Patch classification

A patch classification experiment was conducted following k-fold cross-validation in which a data set was first split into k subsets, and then, in each fold, all except one subset were dedicated to training a classifier model and the remaining one subset was used to evaluate the performance of the model. A training set was usually split further into two sets to keep one as a validation set and estimate the generalization ability of a classifier model and/or convergence of the training process by observing the history of a loss function. As mentioned above, a data set was prepared for three-fold cross-validation so that those images obtained from the same block were not included in both training and test sets to demonstrate how well a classifier model could perform in a practical situation (Fig. 3). Furthermore, some grids of a block were picked out from a training set to construct a validation set.

A batch normalized version^18^ of AlexNet^11^ was adopted to conduct patch classification. The network architecture and some hyperparameters are depicted in Figure 4. Starting from a publicly available model pretrained on the ImageNet dataset,^37^ the classifier model was retrained or fine-tuned on our TEM image/patch set. In the training phase, Gaussian noise was added to a patch, and also, a patch was randomly rotated by either 0, 90, 180, or 270 degrees and flipped horizontally and vertically, before being fed into the classifier model, where the standard deviation of Gaussian noise was randomly chosen in range 0–30. The batch size was set to 32, and 19,200 patches were randomly sampled from the training set in each epoch, so the number of iterations per epoch was 600. To update the model parameters (weights), the Adam optimization algorithm^38^ was used with an initial learning rate of 0.0001 and weight decay of 0.00001. After one epoch training, the validation dataset was fed into the classifier model to calculate the loss, and the learning rate was reduced by a factor of 10 if there was no improvement in the validation loss after five epochs. Cross-entropy loss was used with weights inversely proportional to class frequencies. Thus, the training phase proceeded by alternating between updating the weights and adjusting the learning rate.

Patches obtained from the ENX-resistant strains were treated as a positive class, while patches obtained from the parental strain were treated as a negative class. The performance of the classifier model could be evaluated through sensitivity and specificity defined as follows:

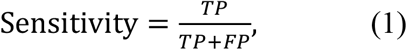

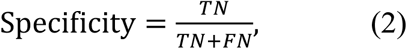

where TP denotes true positives (correctly classified ENX-resistant cell patches), FP denotes false positives (parental cell patches that were classified into the ENX-resistant cell), TN denotes true negatives (correctly classified parental cell patches), and FN denotes false negatives (ENX-resistant cell patches that were classified into the parental cell).

### Visualization of discriminative parts of bacterial cells

Visualization of the characteristic features of both the ENX-resistant and parental cells was attempted using Grad-CAM.^19^ In our approach, inputs to the classifier model were patches, not images. Because it was known which image each patch was extracted from, the class activation map of a patch could be accumulated on its corresponding area within the image. A class activation map was multiplied by the output of the softmax function corresponding to its class, and also, averaged in an area overlapped by some patches. Then, patchwise class activation maps were accumulated onto their originating images to visualize the discriminative parts of the bacterial cells in each image.

### Calculation of Pearson’s correlation coefficients

To investigate which genes were associated with the morphological features, we calculated Pearson’s correlation coefficients between the gene expression information^1^ and the image features extracted from the full connection layer before the last one in the CNN model. We used the expression values of a gene from five strains (ENX-1,2,3,4 and parent) *X* = [*x*_1_, *x*_2_, *x*_3_, *x*_4_ *x*_5_]^*T*^ and also the values of an image feature *Y* = [*y*_1_, *y*_2_, *y*_3_, *y*_4_ *y*_5_]^*T*^ from those five strains, and Pearson’s correlation coefficient r between the level of gene expression and the image feature was defined as

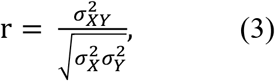

where 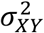 is the covariance of *X* and *Y* and 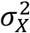 and 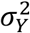 are the variance of *X* and *Y*, respectively. We retained pairs of genes and image features meeting the condition that the absolute correlation coefficient between them was equal to or greater than 0.999. While the image features were extracted as a 4,096-dimensional vector, there were some features whose values were zero for all strains. We removed these features, and consequently, the number of features varied in the cross-validation folds (Table S2). The most frequently appearing genes were highly correlated with image features.

## Acknowledgments

We are grateful to Masashi Yamaguchi at Medical Mycology Research Center, Chiba University, for useful advice on the cryofixation of bacterial cells; Natsue Sakata at RIKEN Center for Biosystems Dynamics Research for technical assistance on single colony isolation of bacterial strains; and Miki Ikebe at Graduate School of Pharmaceutical Sciences, Osaka University, for assistance with the correlation analysis. We are grateful to the technical staff of the Comprehensive Analysis Center of the Institute of Scientific and Industrial Research at Osaka University. This work was performed in part under the Collaborative Research Program of the Institute for Protein Research, Osaka University, CR-19-05.

## Additional information

### Funding

This work was supported in part by MEXT (Ministry of Education, Culture, Sports, Science and Technology of Japan)/JSPS (Japan Society for the Promotion of Science) KAKENHI Grant Numbers 17K08827, 17H06422, 17H03983, 18K19451, 21H03542, by the Naito Foundation, by Takeda Science Foundation, by Network Joint Research Center for Materials and Devices, by Dynamic Alliance for Open Innovation Bridging Human, Environment and Materials, by the Center of Innovation Program and Core Research for Evolutional Science and Technology (JPMJCR20H9) from Japan Science and Technology Agency, JST.

## Contributions

M.H.N., K.A., C.F., and K.N. conceived the experiments; M.H.N., Y.T., K.U., and K.A. performed the LM experiments and morphometric analysis; M.H.N., A.K., Y.T., and A.F. performed the TEM sample preparation and image acquisition; M.H. and K.I. provided assistance with cryofixation; K.A. and A.K. conducted the TEM image classification by deep learning; E.S., T.W., and K.A. performed the calculations of correlation coefficients; M.H.N., K.A., and K.N. wrote the manuscript; T.E., Y.Y., and K.N. supervised the project. All authors reviewed and approved the final manuscript.

## Competing interests

The authors declare no competing interests.

## Notes

### Competing Interest Statement

The authors have declared no competing interest.

## References

1. Suzuki, S., Horinouchi, T. & Furusawa, C. Prediction of antibiotic resistance by gene expression profiles. Nat Commun 5, 5792, doi:10.1038/ncomms6792 (2014).

2. Furusawa, C., Horinouchi, T. & Maeda, T. Toward prediction and control of antibiotic-resistance evolution. Curr Opin Biotechnol 54, 45–49, doi:10.1016/j.copbio.2018.01.026 (2018).

3. Maeda, T. et al. High-throughput laboratory evolution reveals evolutionary constraints in Escherichia coli. Nat Commun 11, 5970, doi:10.1038/s41467-020-19713-w (2020).

4. Giesbrecht, P., Kersten, T., Maidhof, H. & Wecke, J. Staphylococcal cell wall: morphogenesis and fatal variations in the presence of penicillin. Microbiol Mol Biol Rev 62, 1371–1414 (1998).

5. Elliott, T. S., Shelton, A. & Greenwood, D. The response of Escherichia coli to ciprofloxacin and norfloxacin. J Med Microbiol 23, 83–88, doi:10.1099/00222615-23-1-83 (1987).

6. Zhu, Y., Ouyang, Q. & Mao, Y. A deep convolutional neural network approach to single-particle recognition in cryo-electron microscopy. BMC Bioinformatics 18, 348, doi:10.1186/s12859-017-1757-y (2017).

7. Zeng, T., Wu, B. & Ji, S. DeepEM3D: approaching human-level performance on 3D anisotropic EM image segmentation. Bioinformatics 33, 2555–2562, doi:10.1093/bioinformatics/btx188 (2017).

8. Hunter, R. C. & Beveridge, T. J. High-resolution visualization of Pseudomonas aeruginosa PAO1 biofilms by freeze-substitution transmission electron microscopy. J Bacteriol 187, 7619–7630, doi:10.1128/JB.187.22.7619-7630.2005 (2005).

9. Vanhecke, D., Graber, W. & Studer, D. Close-to-native ultrastructural preservation by high pressure freezing. Methods Cell Biol 88, 151–164, doi:10.1016/S0091-679X(08)00409-3 (2008).

10. Kellenberger, E. The ‘Bayer bridges’ confronted with results from improved electron microscopy methods. Mol Microbiol 4, 697–705, doi:10.1111/j.1365-2958.1990.tb00640.x (1990).

11. Krizhevsky, A., Sutskever, I. & Hinton, G. E. ImageNet classification with deep convolutional neural networks. Adv Neural Inf Process Syst, 1097–1105. (2012).

12. Russakovsky, O. et al. ImageNet large scale visual recognition challenge. Int J Comput Vis 115, 211–252, doi:10.1007/s11263-015-0816-y (2015).

13. Fakhry, A., Zeng, T. & Ji, S. Residual deconvolutional networks for brain electron microscopy image segmentation. IEEE Trans Med Imaging 36, 447–456, doi:10.1109/tmi.2016.2613019 (2017).

14. Cireşan, D. C., Giusti, A., Gambardella, L. M. & Schmidhuber, J. Deep neural networks segment neuronal membranes in electron microscopy images. Adv Neural Inf Process Syst, 2843–2851. (2012).

15. Haberl, M. G. et al. CDeep3M—Plug-and-Play cloud-based deep learning for image segmentation. Nat Methods 15, 677–680, doi:10.1038/s41592-018-0106-z (2018).

16. Modarres, M. H. et al. Neural network for nanoscience scanning electron microscope image recognition. Sci Rep 7, 13282, doi:10.1038/s41598-017-13565-z (2017).

17. Weber, G. H., Ophus, C. & Ramakrishnan, L. Automated labeling of electron microscopy images using deep learning. Proc Mach Learn HPC Environ. 26–36. (2018).

18. Ioffe, S., Szegedy, C. Batch Normalization: Accelerating Deep Network Training by Reducing Internal Covariate Shift. 1502.03167v3 [cs.LG] (2015).

19. Selvaraju, R. R. et al. Grad-CAM: Visual explanations from deep networks via gradient-based localization IEEE Int Conf Comput Vis (ICCV), 618–626 (2017).

20. Asmar, A. T. & Collet, J. F. Lpp, the Braun lipoprotein, turns 50-major achievements and remaining issues. FEMS Microbiol Lett 365, doi:10.1093/femsle/fny199 (2018).

21. Dalebroux, Z. D. & Miller, S. I. Salmonellae PhoPQ regulation of the outer membrane to resist innate immunity. Curr Opin Microbiol 17, 106–113, doi:10.1016/j.mib.2013.12.005 (2014).

22. Plumbridge, J. An alternative route for recycling of N-acetylglucosamine from peptidoglycan involves the N-acetylglucosamine phosphotransferase system in Escherichia coli. J Bacteriol 191, 5641–5647, doi:10.1128/JB.00448-09 (2009).

23. Shepherd, M., Sanguinetti, G., Cook, G. M. & Poole, R. K. Compensations for diminished terminal oxidase activity in Escherichia coli: Cytochrome bd-II-mediated respiration and glutamate metabolism. J Biol Chem 285, 18464–18472, doi:10.1074/jbc.M110.118448 (2010).

24. Li, G. W., Burkhardt, D., Gross, C. & Weissman, J. S. Quantifying absolute protein synthesis rates reveals principles underlying allocation of cellular resources. Cell 157, 624–635, doi:10.1016/j.cell.2014.02.033 (2014).

25. Yem, D. W. & Wu, H. C. Physiological characterization of an Escherichia coli mutant altered in the structure of murein lipoprotein. J Bacteriol 133, 1419–1426, doi:10.1128/JB.133.3.1419-1426.1978 (1978).

26. Asmar, A. T. et al. Communication across the bacterial cell envelope depends on the size of the periplasm. PLOS Biol 15, e2004303, doi:10.1371/journal.pbio.2004303 (2017).

27. Hirota, Y., Ryter, A. & Jacob, F. Thermosensitive mutants of E. coli affected in the processes of DNA synthesis and cellular division. Cold Spring Harb Symp Quant Biol 33, 677–693, doi:10.1101/sqb.1968.033.01.077 (1968).

28. Wachi, M. et al. Mutant isolation and molecular cloning of mre genes, which determine cell shape, sensitivity to mecillinam, and amount of penicillin-binding proteins in Escherichia coli. J Bacteriol 169, 4935–4940, doi:10.1128/jb.169.11.4935-4940.1987 (1987).

29. Hews, C. L., Cho, T., Rowley, G. & Raivio, T. L. Maintaining integrity under stress: Envelope stress response regulation of pathogenesis in Gram-negative bacteria. Front Cell Infect Microbiol 9, 313, doi:10.3389/fcimb.2019.00313 (2019).

30. Hengge-Aronis, R. Signal transduction and regulatory mechanisms involved in control of the sigma(S) (RpoS) subunit of RNA polymerase. Microbiol Mol Biol Rev 66, 373–395, table of contents, doi:10.1128/mmbr.66.3.373-395.2002 (2002).

31. Gutierrez, A. et al. Beta-lactam antibiotics promote bacterial mutagenesis via an RpoS-mediated reduction in replication fidelity. Nat Commun 4, 1610, doi:10.1038/ncomms2607 (2013).

32. Gruber, T. M. & Gross, C. A. Multiple sigma subunits and the partitioning of bacterial transcription space. Annu Rev Microbiol 57, 441–466, doi:10.1146/annurev.micro.57.030502.090913 (2003).

33. Keseler, I. M. et al. The EcoCyc database: Reflecting new knowledge about Escherichia coli K-12. Nucleic Acids Res 45, D543–D550, doi:10.1093/nar/gkw1003 (2017).

34. Germond, A. et al. Raman spectral signature reflects transcriptomic features of antibiotic resistance in Escherichia coli. Commun Biol 1, 85, doi:10.1038/s42003-018-0093-8 (2018).

35. Aldred, K. J., Kerns, R. J. & Osheroff, N. Mechanism of quinolone action and resistance. Biochemistry 53, 1565–1574, doi:10.1021/bi5000564 (2014).

36. Mori, E., Furusawa, C., Kajihata, S., Shirai, T. & Shimizu, H. Evaluating (13)C enrichment data of free amino acids for precise metabolic flux analysis. Biotechnol J 6, 1377–1387, doi:10.1002/biot.201000446 (2011).

37. Paszke, A. et al. PyTorch: An imperative style, high-performance deep learning library. Adv Neural Inf Process Syst, 8024–8035 (2019).

38. Kingma, D. P. & Ba, J. Adam: A method for stochastic optimization. 1412.6980 [cs.LG] (2014).

