## Supplemental Figures S1, S2, S3, S4, Tables S1, and S2. for "Identification of bacterial drug-resistant cells by the convolutional neural network in transmission electron microscope images"

### Supplementary Materials

Fig. S1. Sample preparation and data acquisition procedures.

Fig. S2. TEM images of bacterial specimens used for the study.

Fig. S3. Preprocessing of TEM images for cell region extraction.

Fig. S4. Additional Grad-CAM results depicting discriminative parts of cells.

Table S1. Number of patches used for the study.

Table S2. Gene groups correlating highly with image features.

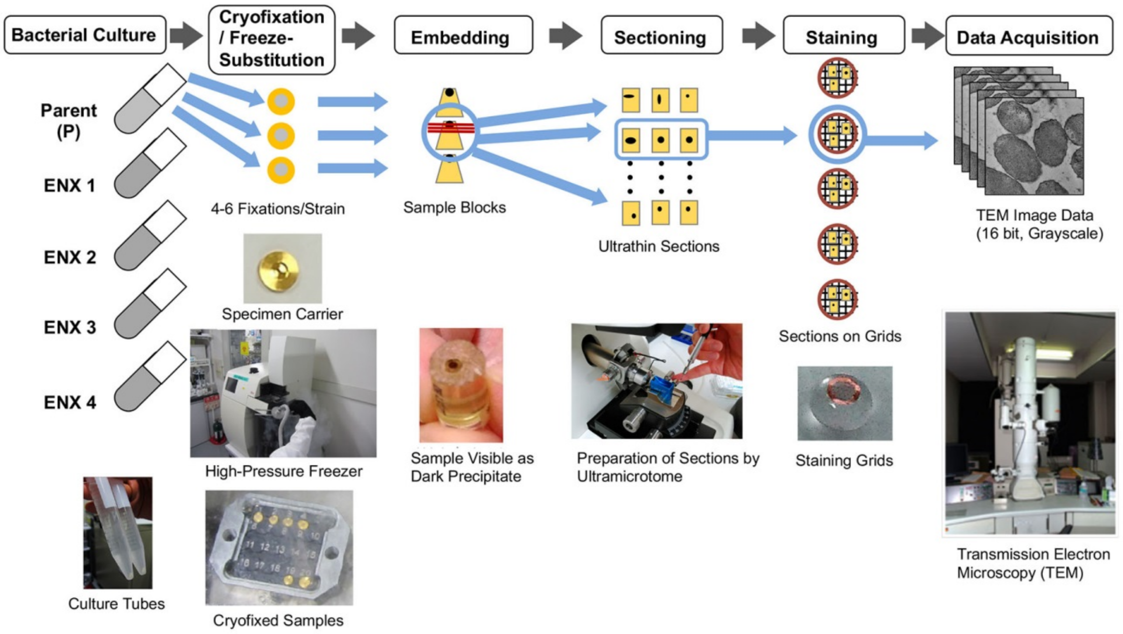


**Fig. S1. Sample preparation and data acquisition procedures.**

The bacterial cell cultures indicated were centrifuged, transferred to a flat specimen carrier, and cryofixed. Cryofixation was conducted several times for each culture. Freeze-substituted specimens were embedded in epoxy resin to make the specimen blocks. Ultra-thin sections were cut from each specimen block and collected on grids. The grids were counterstained and observed under a TEM. Images of bacterial cells were taken and used for analysis.

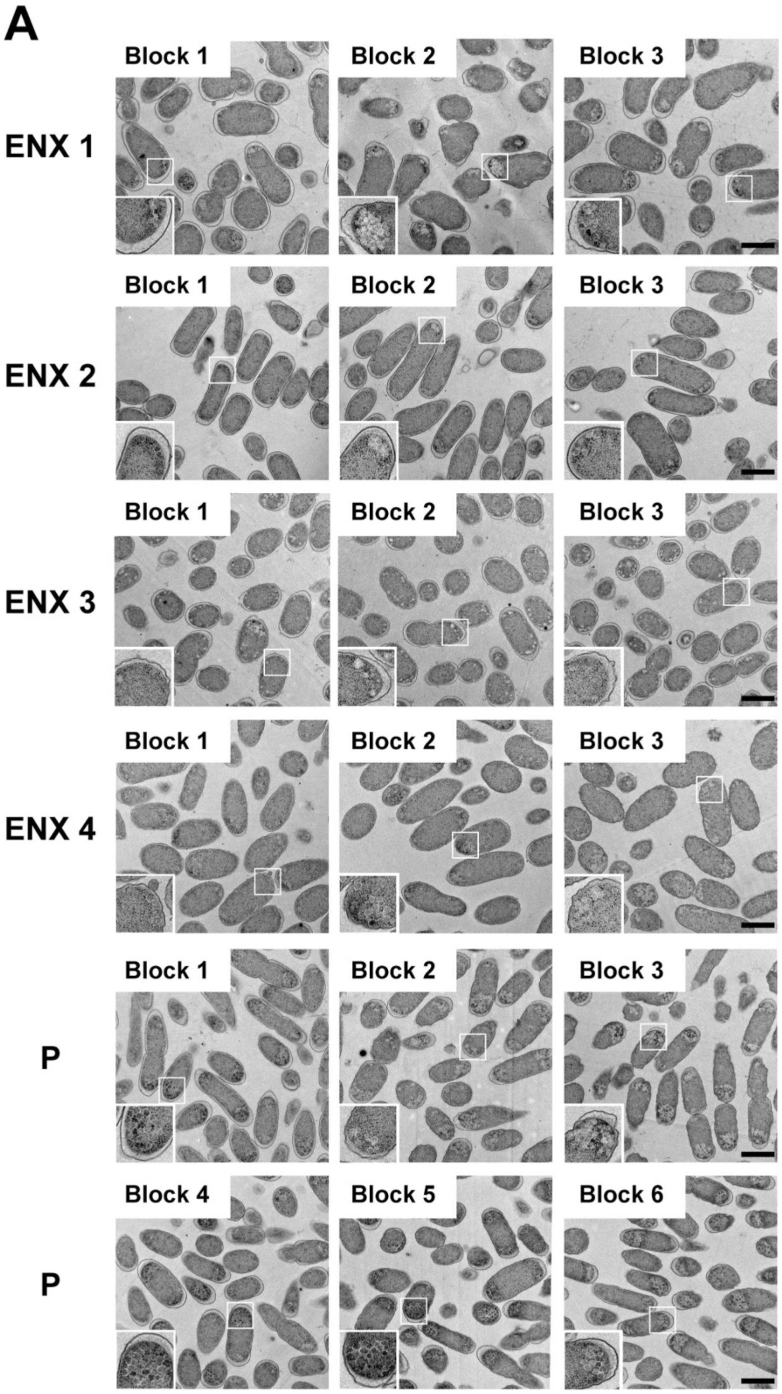
A
Block 1
Block 2
Block 3
ENX 1
ENX 2
ENX 3
ENX 4
P
Block 4
Block 5
Block 6
P

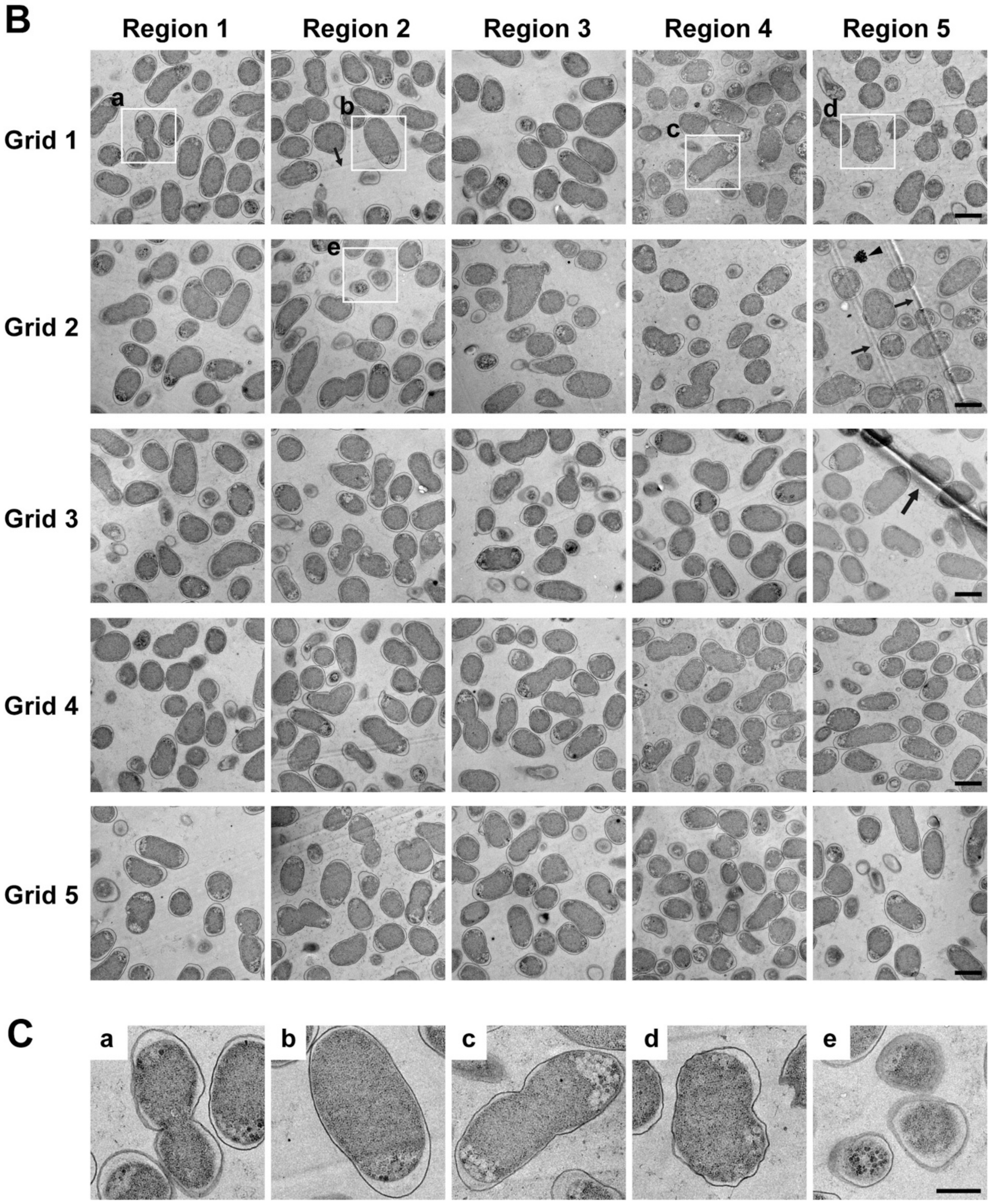


**Fig. S2. TEM images of bacterial specimens used for the study.**

(A) Representative TEM images of 80-nm-thick sections prepared from individual bacterial specimen blocks used for the study are shown. Insets show the magnification of the boxed regions in each micrograph, indicating variations in subcellular appearance, such as the smoothness of membranes, periplasmic space, and electron density of granules, among the specimen blocks. (B) Examples of TEM images of ENX 1 (Block 1) are shown. These were obtained from five different arbitrary regions in the sections on five different grids. (C) Enlarged views of the boxed regions in (B) show examples of large variations in the appearance and shape of cells. (a–d) Vertical cross sections of a dividing cell, single cell, a lower contrast cell with less stained granules, and a cell with irregular shape and rattling outer membrane, respectively. (e) Horizontal cross section of three neighboring cells. Examples of knife marks, wrinkle of a section, and noncellular contamination are indicated by arrows, a large arrow, and an arrowhead, respectively. Scale bars in A and B, 1 μm; in C, 500 nm.

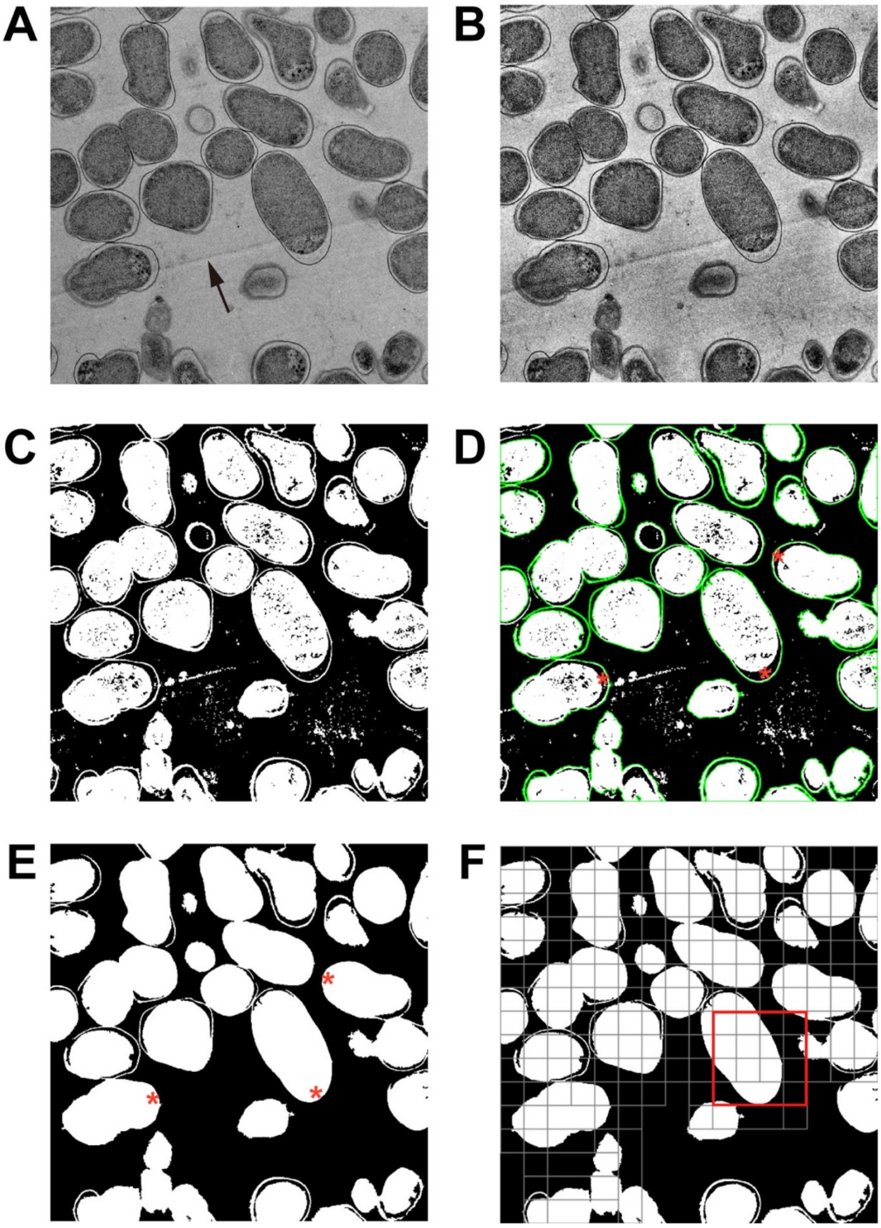


**Fig. S3. Preprocessing of TEM images for cell region extraction.**

(A) A raw TEM image of ENX 1 (the same image from Fig. S2B, Grid 1, Region 2) with uneven intensity levels (after standardization and conversion into 8 bit). Note that the intensity is brighter at upper right of the image. A knife mark is indicated by an arrow. (B) The TEM image after enhancement of the local contrast by CLAHE. (C) A mask image obtained by the Otsu method. (D) Most external contours used to fill holes (i.e., enlarged periplasm, indicated by red asterisks) in cell regions. (E) A mask image refined through hole filling and removal of small areas including a knife mark. (F) Extraction of patches from the TEM images. An example of a 512 × 512 patch containing a portion of cells is indicated by the red square.

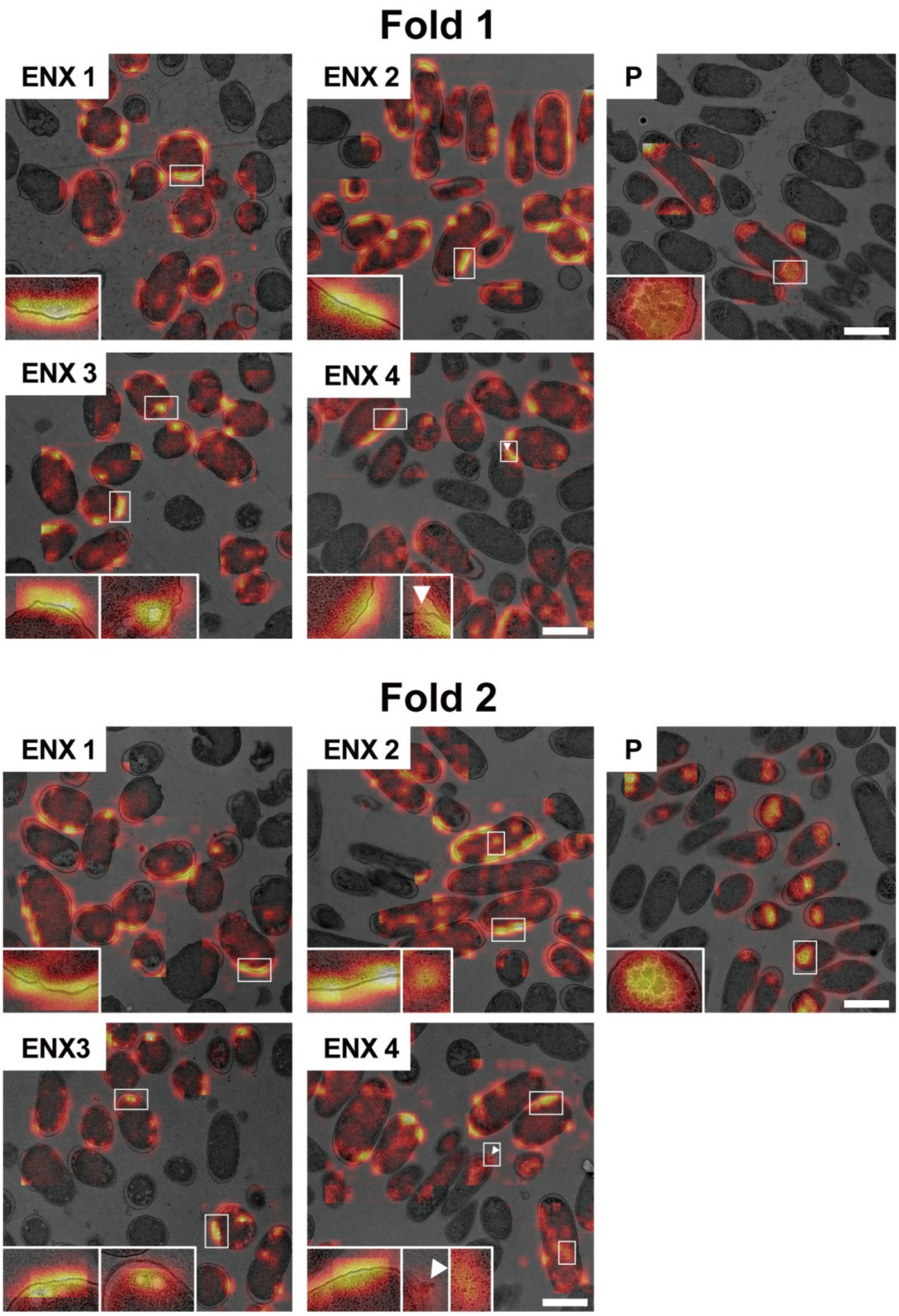


**Fig. S4. Additional Grad-CAM results depicting discriminative parts of cells.**

Besides the Grad-CAM results demonstrated in Figure 6, similar results obtained from the other two folds are shown. Insets depict enlarged views of the boxed regions in each image. White arrowheads indicate bleb-like structures. Scale bars, 1 μm.

**Table S1. Number of patches used for the study[a].**

| Fold | | Training | Validation | Test |
|---|---|---|---|---|
| 1 | Positive | 285,478 | 73,852 | 236,474 |
| | Negative | 231,032 | 69,444 | 128,165 |
| 2 | Positive | 328,161 | 85,702 | 181,941 |
| | Negative | 201,531 | 69,922 | 157,188 |
| 3 | Positive | 331,769 | 86,646 | 177,389 |
| | Negative | 222,399 | 62,954 | 143,288 |

[a]The number of patches extracted from the TEM images and used for training, validation, and test sets, are shown. "Positive" denotes the ENX-resistant strains and "Negative" denotes the parental strain.

**Table S2. Gene groups correlating highly with image features[a].**

| Fold 1 | | | | Fold 2 | | | | Fold 3 | | |
|---|---|---|---|---|---|---|---|---|---|---|
| Gene | Frequency | Mean Value | | Gene | Frequency | Mean Value | | Gene | Frequency | Mean Value |
| *lpp* | 6 | 0.999678 | | *lpp* | 23 | 0.999633 | | *lpp* | 68 | 0.99962 |
| *ybgJ* | 5 | 0.999491 | | *phoP* | 16 | 0.999538 | | *phoP* | 12 | 0.99951 |
| *asnC* | 4 | 0.999403 | | *pspB* | 11 | 0.999549 | | *appB* | 4 | 0.99979 |
| *aroH* | 3 | 0.999182 | | *nagE* | 5 | 0.999376 | | *pdxK* | 2 | 0.99938 |
| *yedQ* | 3 | 0.999350 | | *rrsH* | 3 | 0.999187 | | *yacC* | 1 | 0.99985 |
| *yheO* | 3 | 0.999572 | | *appB* | 3 | 0.999457 | | *hlpA* | 1 | 0.99956 |
| *carA* | 2 | 0.999085 | | *tgt* | 2 | 0.999277 | | *tilS* | 1 | 0.99916 |
| *appB* | 2 | 0.999711 | | *aroH* | 2 | 0.999092 | | *ybdR* | 1 | 0.99903 |
| *ydcW* | 2 | 0.999231 | | *cysC* | 2 | 0.999475 | | *nagE* | 1 | 0.99931 |
| *yeeZ* | 2 | 0.999269 | | *lysA* | 2 | 0.999450 | | *lolE* | 1 | 0.99904 |
| *zipA* | 2 | 0.999522 | | *yheM* | 2 | 0.999968 | | *ycgM* | 1 | 0.99931 |
| *aspT* | 2 | 0.999148 | | *secF* | 1 | 0.999374 | | *dhaL* | 1 | 0.99925 |
| *secF* | 1 | 0.999127 | | *tesB* | 1 | 0.999716 | | *exoX* | 1 | 0.99922 |
| *nagB* | 1 | 0.999407 | | *seqA* | 1 | 0.999245 | | *yedQ* | 1 | 0.99908 |
| *nagE* | 1 | 0.999029 | | *artJ* | 1 | 0.999745 | | *lysA* | 1 | 0.99946 |
| *pepN* | 1 | 0.999764 | | *narX* | 1 | 0.999231 | | *rpoD* | 1 | 0.99918 |
| *hyaB* | 1 | 0.999856 | | *yciN* | 1 | 0.999464 | | *cysG* | 1 | 0.99930 |
| *hyaE* | 1 | 0.999478 | | *ydcW* | 1 | 0.999126 | | *tag* | 1 | 0.99998 |
| *hyaF* | 1 | 0.999504 | | *rspB* | 1 | 0.999444 | | *rpmH* | 1 | 0.99940 |
| *lolE* | 1 | 0.999685 | | *yodA* | 1 | 0.999197 | | *asnC* | 1 | 0.99945 |
| *phoP* | 1 | 0.999533 | | *yeeY* | 1 | 0.999481 | | *cytR* | 1 | 0.99969 |
| *dhaL* | 1 | 0.999690 | | *yeeZ* | 1 | 0.999223 | | | | |
| *rssA* | 1 | 0.999102 | | *yhjR* | 1 | 0.999187 | | | | |
| *oppF* | 1 | 0.999566 | | *ttk* | 1 | 0.999612 | | | | |
| *rspB* | 1 | 0.999647 | | *yieM* | 1 | 0.999241 | | | | |
| *ydgD* | 1 | 0.999316 | | *rplA* | 1 | 0.999379 | | | | |
| *rstA* | 1 | 0.999987 | | *creB* | 1 | 0.999318 | | | | |
| *sanA* | 1 | 0.999183 | | | | | | | | |
| *cysP* | 1 | 0.999125 | | | | | | | | |
| *yfiP* | 1 | 0.999219 | | | | | | | | |
| *clpB* | 1 | 0.999009 | | | | | | | | |
| *rpoD* | 1 | 0.999504 | | | | | | | | |
| *yrbF* | 1 | 0.999078 | | | | | | | | |
| *fdhE* | 1 | 0.999495 | | | | | | | | |

[a]Genes that met the condition that the absolute correlation coefficient was 0.999 or higher, obtained by calculating the Pearson correlation coefficient between the gene

expression information and the image features extracted from the CNN model, are shown. The genes that correlated frequently with the image features are listed in descending order of number of appearances. The number of image features varied in the three folds as 948, 836, and 808, respectively.
